# A Cry For Kelp: Evidence for Polyphenolic Inhibition of Oxford Nanopore Sequencing of Brown Algae

**DOI:** 10.1101/2024.06.26.600698

**Authors:** William S. Pearman, Vanessa Arranz, Jose I. Carvajal, Annabel Whibley, Yusmiati Liau, Katherine Johnson, Rachel Gray, Jackson M. Treece, Neil J. Gemmell, Libby Liggins, Ceridwen I. Fraser, Evelyn L. Jensen, Nicholas J. Green

## Abstract

Genomic resources for macroalgae are increasingly important for conservation and commercial management, however the generation of such resources continues to be hampered by difficulties in the isolation of suitable DNA. Even when DNA has been isolated that otherwise appears high quality, such samples may not perform well during the sequencing process. We here compare Oxford Nanopore long-read sequencing results for three species of macroalgae to those of non-macroalgal species and find that macroalgal samples tend to lead to rapid decline in the number of available sequencing pores resulting in reduced sequencing yield. LC-MS analysis of macroalgal DNA that would be considered suitable for sequencing reveals that DNA derived from dried macroalgae is enriched for polyphenol-DNA adducts – which may lead to sequencing inhibition. Our findings have wide-ranging implications for the generation of genomic resources from macroalgae, for example long read sequencing of dried herbarium specimens, and suggest a need to use fresh tissue wherever possible for genome sequencing.

## Introduction

Contaminants in algal and plant DNA extractions have long been a source of frustration for molecular biologists (Hoarau et al., 2007; Ramakrishnan et al., 2017; Wilson et al., 2016). As whole genome sequencing (WGS) increasingly replaces more traditional genetic approaches, there is a corresponding need for a shift from extraction methods that yield PCR-grade DNA to WGS-grade DNA extractions (De La Cerda et al., 2023; Jones et al., 2021). Notably, some WGS approaches, such as Oxford Nanopore Technologies (ONT) sequencing are particularly sensitive to DNA quality (Pinzauti et al., 2022), since single molecules of DNA must unwind and pass through the nanopore and contaminants in the library may block pores. Thus, DNA that is both high molecular weight and of high ‘quality’ (as judged by UV absorbance ratios, concentration, and electrophoresis) is recommended for nanopore WGS. Despite this, three separate laboratories involved in this research have found consistent difficulties in generating ONT sequencing (hereafter nanopore sequencing) data from three species of brown algae (*Ecklonia radiata, Durvillaea antarctica, Eisenia cockeri*), despite high-quality DNA input, as measured by UV 260/280 and 260/230 absorbance ratios, electrophoresis (Agilent Tape Station or gel electrophoresis), and concentration. Although much focus has been on removing polysaccharide contaminants such as alginates (e.g., Wilson et al. 2016), which often reduces DNA yields, another well-known source of sequencing inhibition is the presence of polyphenols. These latter compounds can oxidise and lead to the formation of DNA-quinone adducts prior to and during DNA extraction (Heikrujam et al., 2020). These adducts may be at such low levels that they are not reliably detected using traditional quality metrics but may still cause problems in WGS.

We noted during numerous sequencing attempts undertaken over several years that our nanopore libraries constructed using brown algal DNA consistently blocked flow cells quickly, and that the average read lengths of our libraries were consistently much shorter than anticipated based on the molecular weight of the input libraries. We hypothesized that, despite employing conventional techniques to suppress such adduct formation during DNA extraction, polyphenol-DNA adducts within our samples could explain the rapid pore blockage that we observed (De La Cerda et al., 2023; Mgwatyu et al., 2022).

Macroalgae are well known for having a high abundance of polyphenols (up to 20% of dry weight; Aminina et al., 2020); these can oxidise into o-quinones, many of which are capable of irreversibly covalently binding to DNA (Heikrujam et al., 2020). Our DNA extractions were conducted using methods optimized to minimise generation and interference by polyphenols, such as the inclusion of high levels of antioxidants and polyphenol sequestering compound polyvinyl-pyrrolidone (PVP). Nevertheless, if adduct formation occurs prior to DNA extraction (e.g., the DNA is natively modified within the cell, or during storage), then such optimized protocols may be for nought.

In this paper we employ both bioinformatics and chemistry approaches to test the hypothesis that polyphenol-derived DNA adducts may interfere with Nanopore DNA sequencing in brown algae, despite using DNA extraction methods aimed to prevent this. Firstly, we expected that macroalgae would tend to promote pore blockages and thus lead to faster pore ‘die off’ than non-macroalgal samples. Secondly, we hypothesized that chemical characterization of the DNA extracts would reveal the presence of DNA-polyphenol adducts. Finally, given our theory that polyphenol-DNA adducts can form prior to extraction, we also expected that DNA derived from silica-dried macroalgae would have higher abundances of these modified bases than fresh tissue, due to cell rupture during storage and/or desiccation.

## Methods

### DNA Extraction, Sequencing, & Basecalling

#### Eisenia cockeri

Tissue was preserved dried on silica beads or stored in RNALater. DNA was extracted from ∼2mm^2^ pieces of tissue using a Qiagen Plant Pro Kit. Tissue was homogenized for 2 minutes in the provided tissue disruption tubes using a Spex Geno/Grinder and incubated overnight at 56 °C, after which it was treated with RNase A (Qiagen). An additional step was carried out following RNase treatment where 100ul of isopropanol was added and incubated at 56°C before the addition of the CD2 solution. Extracted DNA was cleaned up using a Qiagen DNeasy PowerClean CleanUp Kit. The DNA was processed into Nanopore libraries using the SQK-LSK114 kit. The DNA from silica dried tissue was sequenced using two MinION R10.4.1 flowcells, while the RNALater tissue was sequenced using a single run of a PromethION P24 R10.4.1 flow cell at the Newcastle University Genomics Core Facility. Sequencing data was basecalled with Dorado v0.7.0 with the dna_r10.4.1_e8.2_400bps_hac@v4.1.0 or dna_r10.4.1_e8.2_400bps_hac@v4.3.0 models.

#### Ecklonia radiata

Ethanol (95%) or RNALater preserved *Ecklonia* tissue from the young lamina was purified by first grinding with sterile quartz sand and a pestle in a CTAB buffer (2% CTAB, 0.1 M Tris-HCl pH 8.0, 1.4 M NaCl, 20 mM EDTA, 1% PVP, 1% proteinase K), followed by a 2 hour 56°C incubation and a chloroform:isoamyl (24:1) extraction. DNA pellets were washed with 10mL of 70% ethanol and eluted in 500µL of TE buffer pH8. DNA was then RNAse A treated and cleaned using 2X volumes of Ampure XP beads. DNA was sequenced on Oxford Nanopore R9/R10 flowcells using the LSK112/LSK114 kits. Sequencing data was basecalled with Dorado v0.7.0 with the dna_r9.4.1_e8_hac@v3.3 or dna_r10.4.1_e8.2_400bps_hac@v4.2.0 models.

#### Durvillaea antarctica

We utilized Nanopore sequencing data used to generate a draft genome assembly for *D. antarctica* (Fraser et al., 2022). This DNA was extracted from dried tissue (primarily silica drying or freeze drying) using a CTAB protocol (Cremen et al., 2016) which had been modified to include 2% PVP40K and 2% sodium metabisulfite. The DNA was then secondarily purified using the Qiagen DNEasy PowerClean Pro Cleanup Kit following manufacturer’s instructions. A total of six Oxford Nanopore R9.4.1 flow cells were used with the LSK109 sequencing kit to generate the sequence data. In one instance, the DNA sample was cleaned up once more with the Circulomics XS kit to increase read lengths. Prior to sequencing, DNA was confirmed to be of high quality through use of UV absorbance ratios (260/230 of 2.0-2.20 and 260/280 of 1.80-2.0) and high molecular weight either via gel electrophoresis or size selective precipitation via Circulomics XS kit. Sequencing data was basecalled with Dorado v0.7.0 with the dna_r9.4.1_e8_hac@v3.3 model.

#### Vitis vinifera

DNA was extracted from fresh young leaves of *Vitis vinifera* cv. ‘Sauvignon Blanc’, clone UCD1 (FPMS1) using the Nucleomag plant DNA kit (Macherey-Nagel, Düren, Germany), which is based on CTAB extraction protocol, with purification step automated using an Eppendorf EpMotion 5075 liquid-handling robot. DNA concentration was measured using the Qubit broad range kit on a Qubit Flex instrument and purity was determined using a nanodrop 8000 (both from Thermo Fisher Scientific, Waltham, MA, USA).

Library preparation for Nanopore sequencing was performed using the ligation sequencing kit (SQK-LSK114) following the manufacturer’s protocol and sequenced using R10.4.1 flow cells on an ONT PromethION P24 instrument at Bragato Research Institute. Sequencing data was basecalled with Dorado v0.7.0 with the dna_r10.4.1_e8.2_400bps_hac@v4.2.0 model.

#### Additional nanopore data

Additional Nanopore data were retrieved from the ONT Open Data repository. We used *Drosophila melanogaster* data from (Kim et al., 2021) specifically runID 3f6ceb2e-0fb5-4f1c-b4d9-1ecc5c338172, basecalled with dorado v 0.7.0 with the dna_r10.4.1_e8.2_400bps_hac@v4.1.0 model. We used human data retrieved from the Oxford Nanopore Technologies Genome in a bottle project – specifically runID 2264ba8c-03ef-4a79-ab43-bf6f4f18a6f2, base called with dorado v 0.7.0 with the dna_r10.4.1_e8.2_400bps_hac@v4.2.0 model.

### LC-MS Analysis

For LC-MS analysis, a recently developed protocol optimized for DNA extraction (Pearman et al., 2023) was used. DNA was extracted from ∼200mg of chicken breast muscle tissue or ∼100mg of silica dried *Durvillaea antarctica tissue* (stored for 1 month on silica at room temperature). Simultaneously, DNA was also extracted from ∼200mg of flash-frozen fresh *D. antarctica*, frozen with liquid nitrogen and ground to a fine powder. LC-MS analysis also included a purchased salmon sperm DNA control (sonicated salmon sperm DNA - Agilent #201190).

For each sample, DNA was purified with the Qiagen PowerClean Pro kit and eluted with water, except for the salmon sperm DNA which was already suspended in water. Prior to hydrolysis DNA quality was confirmed for the chicken and both types of algal DNA through measurement of 260/280 and 260/230 UV absorbance ratios (chicken, fresh algae, and dried algae had 260/280 ratios of 1.9, 2.2, and 1.8 respectively and 260/230 ratios of 2.1, 2.05, and 2.1 respectively). For each sample, 3μg of DNA (measured using the Qubit Broad Range kit) was hydrolysed to release the constituent nucleobases following (Lowenthal et al., 2019; Shibayama et al., 2016) by hydrolysis using 400μL (four volumes) of neat formic acid at 140°C for 6 hours. Following hydrolysis, the solutions were dried to a pellet at 45°C, and then resuspended in 30μL of HPLC grade water.

LC-MS analyses of products were conducted on a Shimadzu system comprising an SPD-20A UV detector and LCMS-2020. Samples were eluted from an Atlantis T3 column (3μ, 3×150 mm), column temperature 40°C, with LC-MS grade water containing 0.01% formic acid (Solvent A) and acetonitrile (Solvent B) as solvents, at a flow rate of 0.5 mL/min, with the following gradient: 0 – 6 min, 0 – 1% B; 6 – 22.5 min, 1 – 10% B, 22.5 – 32.5 min, 10 – 100% B, 32.5 – 48.5 min, 100% B; 48.5 – 49.5 min, 100 – 1% B; 49.5 – 52.5 min, 1% B. Total Ion Count was measured between m/z 104 and 1200 in electrospray ionisation (ESI) positive and negative modes. UV was monitored at 260 and 280 nm. Flow rate of nebulising gas was 3 L/min, desolvation line temperature was 250°C, heat block temperature was 400°C, and drying gas flow rate was 15 L/min.

Predicted m/z distributions for polyphenol-derived DNA adducts were produced by summing the masses of each nucleobase and each positive polyphenolic ion previously observed in *Durvillaea antarctica* (Olate-Gallegos et al., 2019), and deducting two to account for the oxidation. Although crude, this approach is intended to give a rough estimation of the distribution of adduct sizes.

### Bioinformatic analysis

We categorized the reasons for nanopore reads ending into two groups – blockage (signal_negative and unblock_mux_change) and non-blockage (signal positive, mux_change). In some instances, flow cells had terminated sequencing early – either the result of extremely poor sequencing, or, in the case of *Drosophila*, to load wash and load additional libraries. These discrepancies make comparisons across runs inherently difficult, nevertheless we present the proportion of pores available for sequencing over the course of each sequencing run, as a proportion of the maximum available throughout the run.

Following basecalling, we identified the terminal base and quality score for each read, and removed those with a Q-score < 8, corresponding to probability of correctly being called of 0.85. Although terminal bases are known to typically have lower Q-scores than the rest of the read (Delahaye & Nicolas, 2021), we would expect that these patterns should equally affect macroalgal and non-macroalgal reads and thus biases in terminal base patterns may be informative for understanding reasons for macroalgal inhibition of nanopore sequencing.

## Results

### Nanopore Sequencing

We observed that macroalgal sequencing runs typically declined quickly in the number of pores available for sequencing, while non-macroalgal sequencing had greater preservation of pores throughout the sequencing run (Fig. 1a). One notable exception was sequencing of RNALater stored *Eisenia* tissue which had pore preservation closer in line with non-macroalgal samples. Secondly, we found reads which terminated because of a blockage were more likely to terminate at a purine (Fig. 1b), while ‘complete’ reads (reads which successfully navigated the pore in their entirety) were far more likely to terminate at pyrimidines. These purine/pyrimidine biases were typical across most sample types except for the human dataset.

**Figure 1:**
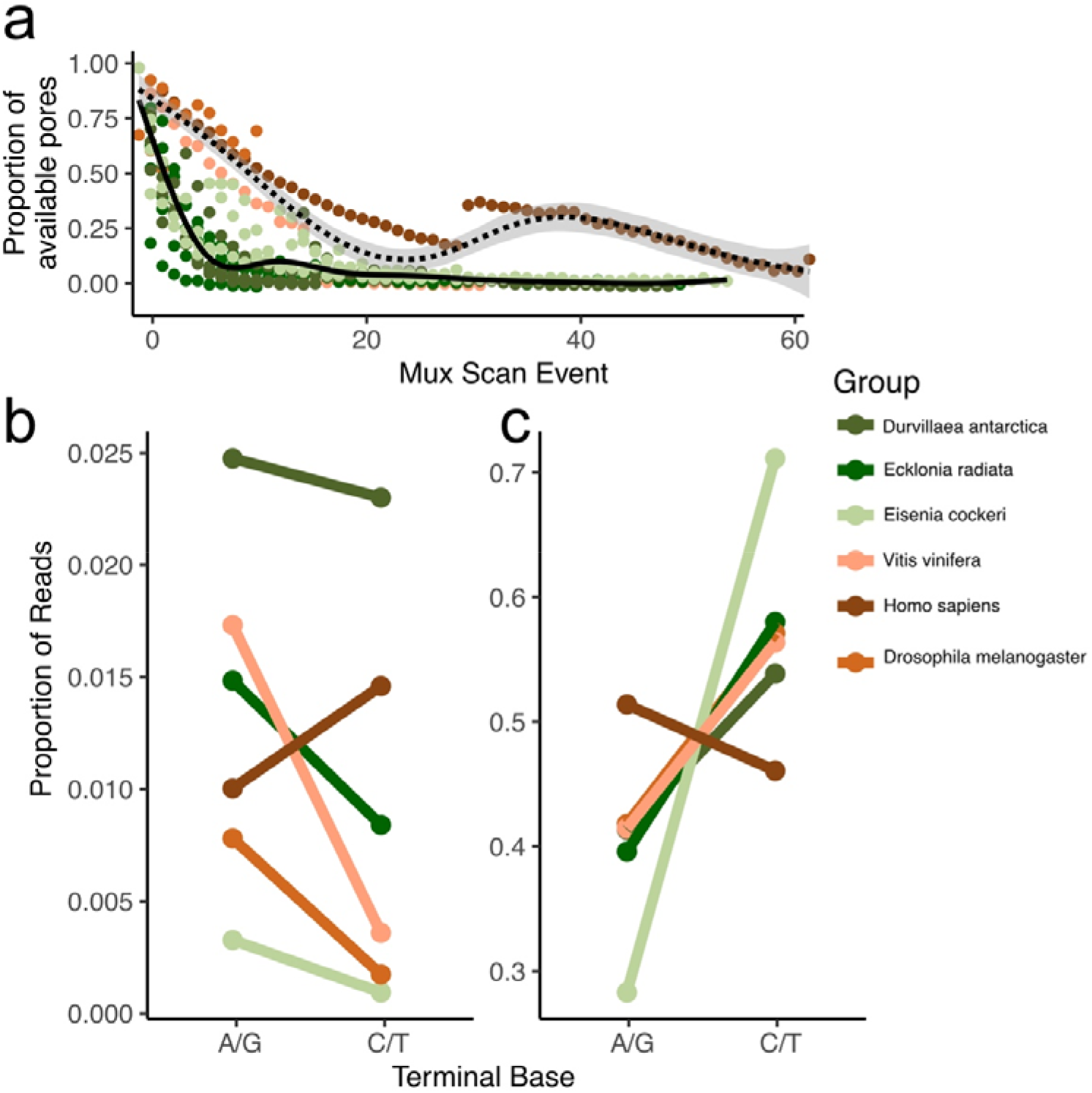
a) Proportion of pores (of maximum available, to account for different numbers of initial pores in MinION and PromethION flow cells) available for sequencing, dashed line indicates GAM smoothing across non-macroalgal samples, solid line indicates smoothing for macroalgal samples b,c) Frequency of reads terminating with A/G vs C/T for reads which terminated sequencing because of a blockage (b) and reads which terminated due to completing sequencing the read (c).

**Figure 2:**
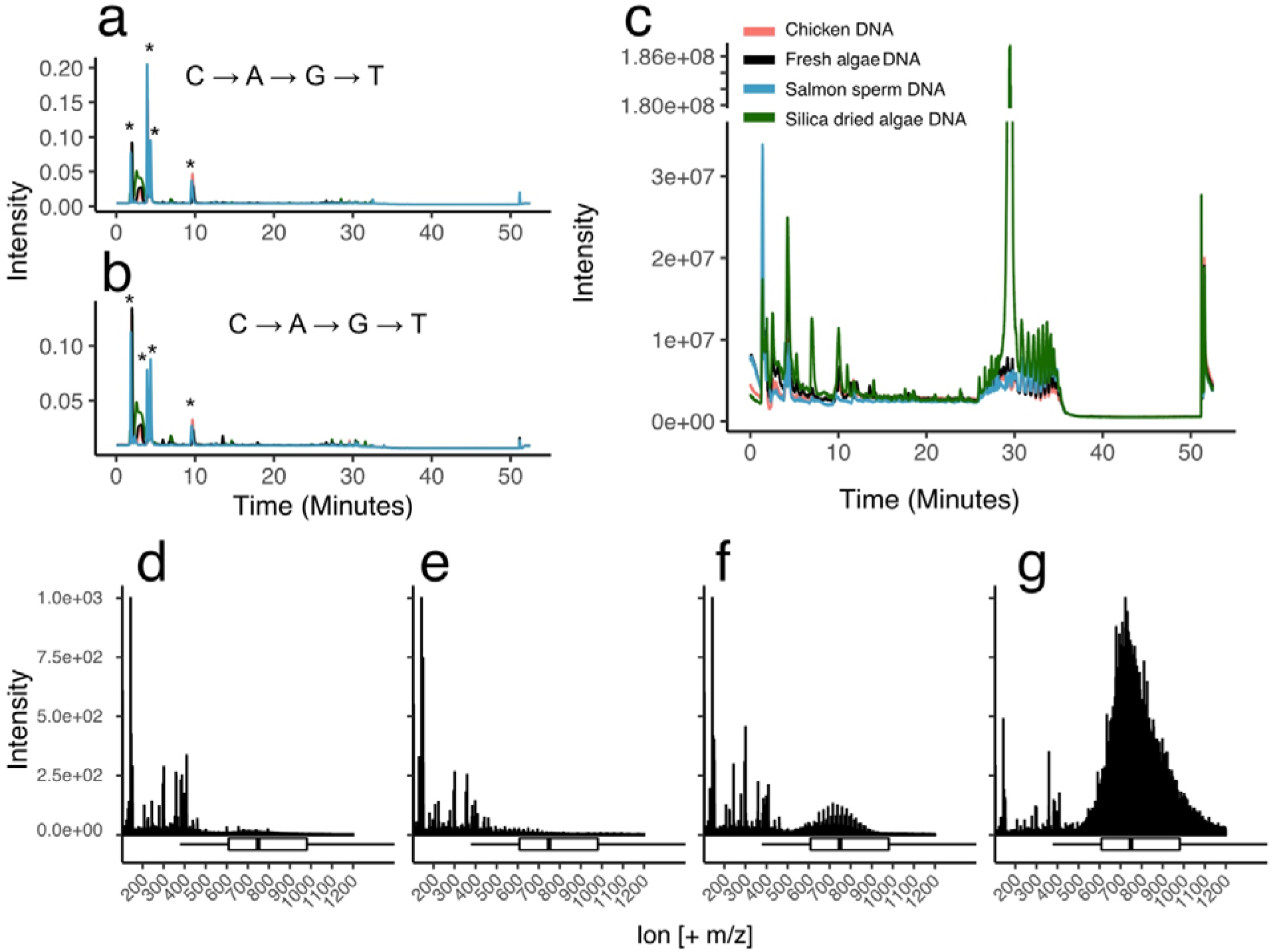
a) Overlaid UV chromatogram at 280 nm of hydrolysed DNA for each DNA extract, stars represent the absorbance peaks for each base, confirmed through analysis of extracted ion chromatograms. C through T is the order of the DNA bases relative to the stars. b) Same as a, except at 260nm. c) Overlaid total ion chromatograms of all DNA samples, note the axis break in intensity for silica dried algal DNA. d-g) Combined mass spectra between 29 and 30 minutes for each sample (salmon, chicken, fresh algae, dry algae respectively). Boxplots below d-g are the hypothesized distribution of polyphenol derived DNA adducts for *Durvillaea*, calculated as described in the Methods section.

**Figure 3:**
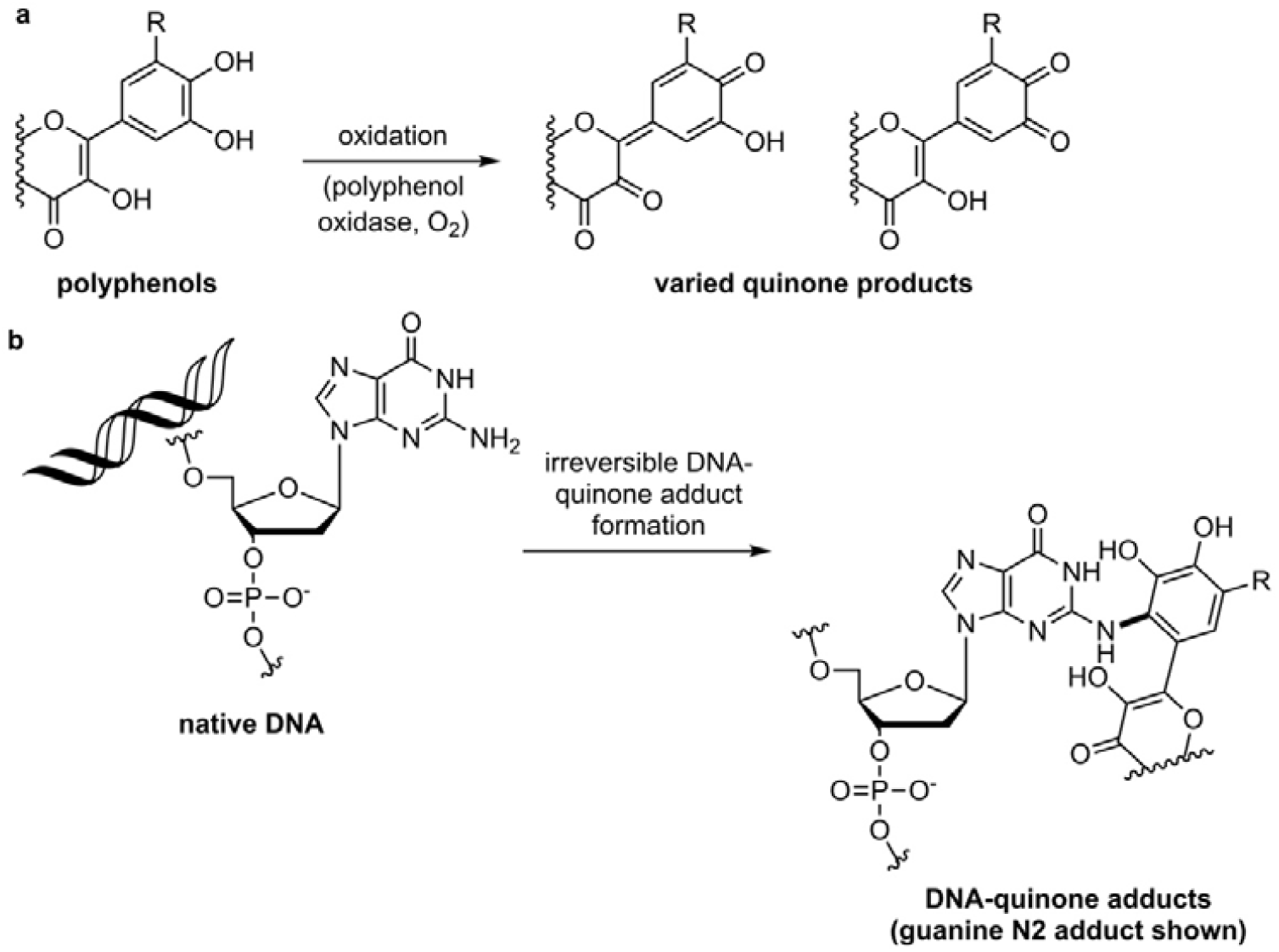
Proposed mechanism for accumulation of DNA-polyphenol adducts. a) Example flavanol polyphenol undergoing oxidation (b), reaction with guanine DNA base forming a flavanol derived o-quinone DNA adduct (new covalent bond in bold).

### LC-MS

We observed the presence of four major peaks by UV at 280 nm, which aligned with the position of the extracted ion chromatograms of the DNA bases cytosine, adenine, guanine, and thymine. Examination of the total ion chromatogram (TIC) of negatively ionized samples revealed no differences between any samples, however the TIC of positively ionized samples revealed a singular major difference between the dried algal sample and all others – an extremely large peak at 29.5 minutes. Analysis of the ion composition of our data between 29 and 30 minutes revealed this peak was dominated by high molecular weight products aligning with the hypothesized distribution of polyphenol derived DNA base adducts. A distribution was also observed in the fresh algal DNA over this retention time, however it was substantially smaller than in dried algal DNA and was not apparent on the TIC.

## Discussion

Despite increasing interest in the production of genomic resources for brown algae, our three groups have all independently encountered extensive difficulties in producing high quality Nanopore sequencing data for these algae. Given the well-established role of polyphenols in interfering in DNA extractions (Hoarau et al., 2007; Jones et al., 2021), and the known high abundance of polyphenolic compounds in macroalgae (Aminina et al., 2020), we suggest based on two independent lines of evidence that polyphenol-DNA adduct formation, dependent on sample processing methods, may underlie problematic next generation sequencing results. First, we suggest that, based on the available Nanopore data and additional LC-MS data generated, DNA-orthoquinone adducts can form during sample drying prior to DNA extraction. We demonstrate here that DNA extracted from silica-desiccated tissue of *D. antarctica* possess a distribution of high molecular weight compounds aligning with hypothesized polyphenol-derived DNA adducts. Conversely, DNA extracted in a similar fashion from either fresh tissue, or from animal tissue, possess fewer (in the case of fresh tissue) or none of these compounds.

Notably, in all instances of both Nanopore sequencing and LC-MS, our extracted DNA would be considered ‘high quality’ by traditional metrics of assessment (UV absorbance ratios, Qubit, gel electrophoresis). We suggest that this is likely because polyphenol modified bases make up a very small proportion of the overall base composition of our DNA extractions, and because polyphenols frequently share a UV absorbance spectrum with DNA. For many approaches, such modified bases may not represent a major source of inhibition, especially those which involve the use of PCR amplification - as sufficient levels of unmodified template DNA are likely present to minimize these issues. However, processes which directly utilize native DNA, such as nanopore whole genome sequencing, may be more susceptible to these issues. For example, if even 0.01% of bases are modified, this would correspond to 10,000 bases per megabase – given that sequencing yields are, ideally, in the gigabase range, the potential for accumulation of modified bases blocking or damaging the pores could lead to reduced sequencing output is significant.

When analysing formic acid-hydrolysed DNA by LC-MS, we did not observe intense peaks (compared to nucleobase signals) by UV-detection or negative mode ionisation for putative adducts, consistent with the idea that adduct formation is relatively minor. However, proposed DNA-quinone adducts were clearly identified under positive ionization mode, demonstrating the utility of LC-MS as a measure of DNA quality in this context. Previous research has used positive ionization to detect quinone and bisphenol adducts (Zhao et al., 2018), with some orthoquinones being better detected under positive ionization (Es-Safi et al., 2007). Unoxidised polyphenols in *Durvillaea* have been well identified using both negative and positive ionization (Olate-Gallegos et al., 2019). In addition, the formic acid treatment of DNA employed herein may promote the formylation of the hydroxyl groups of polyphenols, reducing their sensitivity to negative ionization, while simultaneously increasing the sensitivity to positive ionization.

Principally, we saw strong LC-MS based evidence for polyphenol derived DNA adducts within silica dried macroalgal tissue, with substantially less of these putative adducts in fresh macroalgal DNA. Indeed, given their substantially lower abundance in fresh tissue we suggest silica or dry storage of tissue may reduce the amenability of brown algae to nanopore sequencing. This is also somewhat supported by the improved sequencing results of *Eisenia* tissue stored in RNALater rather than on silica gel.

### Potential mechanisms

Because our DNA extraction protocols are designed to reduce polyphenol effects on DNA (e.g., use of antioxidant and PVP), we suggest that the majority of DNA degradation arises prior to extraction, most likely during sample storage. Our results align with previous research which has suggested that silica desiccation may be problematic due to water stress and cell damage promoting DNA damage (Akinnagbe et al., 2011; Bainard et al., 2010). In essence, because the desiccation process ruptures cellular compartments, polyphenol oxidase, polyphenols, and nucleic acids are liberated into a common cellular medium – allowing for DNA damage to occur during the drying process (Savolainen et al., 1995). Therefore, based on our data and those of Savolainen et al. (1995), we hypothesize that such storage of samples leads to varying degrees of cell rupture, promoting the release of polyphenol oxidase from chloroplasts and vesicles (Jukanti, 2017), facilitating the oxidation of polyphenols to ortho-quinones. Ortho-quinones are highly reactive species that undergo reaction with DNA, the products of which we suggest are the principal reason for poor nanopore sequencing of brown algae. Therefore, sample storage approaches which preserve cellular integrity, or maintain temperatures which prevent these reactions from occurring, are likely the best steps forward in improving Nanopore sequencing yields.

Finally, the observation of a bias in kelp samples towards blockage-terminated sequencing reads ending with a purine, vs complete sequencing reads ending with a pyrimidine, is noteworthy. Although purines are known to be far more reactive than pyrimidines towards o-quinones (Cavalieri et al., 2002; Wang et al., 2021; Xiong et al., 2022), the origins of this bias require further investigation, since the position of a blockage relative to a nucleobase-quinone adduct, and sequence and structure dependent biases for o-quinone-adduct site formation, remain unknown. One explanation consistent with our data would be the tendency of o-quinone-adducts to form in purine-rich stretches of DNA, resulting in blockages during sequencing of that area. For example, G-quadruplexes are guanine-rich secondary DNA structures which are more readily cross-linked by polyphenol derivatives (Yuan et al., 2013). Such polyphenol derivative compounds can also cross-link double-stranded DNA (Bai et al., 2010) which may also explain the rapid flow cell blockage observed with macroalgal samples.

## Conclusion

We suggest that polyphenolic compounds in brown algae interfere with Nanopore sequencing. Thus, we suggest that tissue storage conditions are important and should in future be considered in the context of whole genome sequencing methods, especially for brown algal samples. Specifically, storage over silica gel should be avoided. Our preliminary findings have significant implications not just in analysis of contemporary macroalgae samples, but also potentially for historic herbarium specimens which may not be amenable to long-read DNA sequencing as a result of DNA adduct formation, in addition to DNA fragmentation from sample age. Additionally, given the ecological and economic importance of macroalgae and the relative paucity of available genomic resources for both commercial and conservation management, there is a pressing need to resolve issues relating to DNA sequencing. Further work is required for quantitative comparisons between samples derived from different processing methods, and our results provide a basis for which LC-MS analysis can be used in this application.

## Acknowledgements

The *Eisenia* data was supported by the National Environment Research Council grants NE/S011692/2 and 2022GCBCKELPER2. The *Ecklonia* work was supported by the New Zealand Ministry for Business Innovation and Employment (MBIE) Strategic Science Investment Fund (SSIF) platform Genomics Aotearoa “High Quality Genomes and Population Genomics” project. The *Durvillaea* sequencing was supported by the Royal Society of New Zealand via a Rutherford Discovery Fellowship to C.I.F. (RDF-UOO1803). The *Vitis vinifera* data were generated by Jessica Rivera-Perez and supported by funding from New Zealand Ministry of Primary Industries and from New Zealand Winegrowers.

